# SurvBenchmark: comprehensive benchmarking study of survival analysis methods using both omics data and clinical data

**DOI:** 10.1101/2021.07.11.451967

**Authors:** Yunwei Zhang, Germaine Wong, Graham Mann, Samuel Muller, Jean Y.H. Yang

**Author notes:** Equal contribution (Samuel Muller, Jean Y.H. Yang).

## Abstract

Survival analysis is a branch of statistics that deals with both, the tracking of time and of the survival status simultaneously as the dependent response. Current comparisons of survival model performance mostly center on clinical data with classic statistical survival models, with prediction accuracy often serving as the sole metric of model performance. Moreover, survival analysis approaches for censored omics data have not been thoroughly investigated. The common approach is to binarise the survival time and perform a classification analysis.

Here, we develop a benchmarking framework, SurvBenchmark, that evaluates a diverse collection of survival models for both clinical and omics datasets. SurvBenchmark not only focuses on classical approaches such as the Cox model, but it also evaluates state-of-art machine learning survival models. All approaches were assessed using multiple performance metrics, these include model predictability, stability, flexibility and computational issues. Our systematic comparison framework with over 320 comparisons (20 methods over 16 datasets) shows that the performances of survival models vary in practice over real-world datasets and over the choice of the evaluation metric. In particular, we highlight that using multiple performance metrics is critical in providing a balanced assessment of various models. The results in our study will provide practical guidelines for translational scientists and clinicians, as well as define possible areas of investigation in both survival technique and benchmarking strategies.

## 1. Background

Survival models are statistical models designed for data that have censored observations, that is time-to-event data, which are ubiquitous, including in health, biology, economics, and engineering. We will follow the terminology in survival analysis and capture the event of interest through a ‘status’ variable, which is typically a binary class outcome; we call the waiting time to this status event the ‘survival’ time, which is either measured as a continuous or discrete time. This class of models has wide applicability well beyond the clinical and omics datasets considered in this article. Survival models target both outcomes: status and time-to-event, whereas neither regression analysis on time nor classification analysis on status explain this bivariate outcome information [1].

Numerous survival models have been developed over the last decades. There are many studies in the literature that give a good overview [2]. However, there is a lack of studies from a practical viewpoint, that is a lack of extensive real-world dataset comparisons, particularly in the biomedical field. This motivates us to develop a benchmarking framework for the diverse clinical and omics survival data in health, to provide a better understanding of such models, and to inform on how to guide decision making. To better understand what has been done, we perform an exhaustive search for various types of available survival analysis methods together with performance evaluations for different types of datasets.

Among the comparison studies that include real-world datasets in health, we found that they typically have specific focus such as on a certain disease (e.g. colon cancer), or on a certain data platform (e.g. omics or clinical). For example, [3] and [4] conduct reviews on classical survival models such as the Kaplan-Meier (KM) method and the Cox Proportional Hazards (CoxPH) model with a focus on clinical data with an induce anesthesia state and a specific colon cancer type, respectively. [5] apply the penalised Cox model, survival support vector machine (SVM), random survival forest (RSF), and Cox boosting models on large genomic data. To date, no systematic review encompasses datasets obtained from multiple disease types and includes both, clinical categorical and clinical continuous data, as well as large omics data. This necessitates the development of a benchmarking framework that will provide a better understanding of the practical impact of different survival models.

With the emergence of different modelling approaches from various disciplines many of these recent comparison studies have limited their focus on either within classical models (KM method, CoxPH model) or within modern machine learning (ML) methods. Recently, a comprehensive survey article by [6] compares three categories of statistical survival and ML methods with a focus on theoretical mathematical detail. However, their study does not discuss practical implications of the various methods and no comparison of performance using real-world datasets. There is a need for better guidance on what data analysis strategy to use.

A recent exception is the benchmark study by [6], this valuable contribution includes both real-world clinical and omics datasets and analyses these with classical regression and modern ML methods with particular focus on the impact of considering the multi-omics structure to the survival model predictability. However, this study includes cancer diseases only and datasets are obtained from ‘The Cancer Genome Atlas’ (TCGA) only. There remains a pressing need to look into more diverse and thus more heterogenous datasets coming from multiple databases to benchmark the survival model performances from more diverse aspects.

To this end, we develop a benchmarking framework: SurvBenchmark that considers multiple aspects with several evaluation metrics on a large collection of real-world health and biomedical datasets to guide method selection and new method development.

## 2. Survival models and their evaluation

In this section, we briefly introduce survival models together with their evaluation metrics. Survival models can deal with data that explain censored observations with a bivariate outcome variable, consisting of ‘time’ (the minimum of ‘time-to-event’ and ‘censoring time’) and ‘event’ (binary: “class 1”= “event did occur”, “class 0”= “otherwise”). There are two key features of such censored survival objects. Firstly, the class label “0” (referring to a “no-event”) does not mean an event class labelled as “0”, instead, it represents that the event-outcome is “unknown”, or, using the technical term, “censored”. Secondly, an additional tracking time measurement is included as part of the response.

There are two main branches of survival models: classical statistical survival models, which include parametric, nonparametric and semi-parametric models; and modern ML survival models, which include ensemble based methods and state-of-the art deep learning based approaches. Both sets of models are briefly reviewed in the following sections.

### 2.1 Classical survival models

The Cox Proportional Hazards (CoxPH) model [7] is the most widely used classical survival model. CoxPH works on the hazard function, which is given by

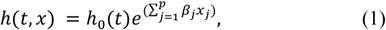

where *x* = (*x*_1_, *x*_2_, …, *x*_*p*_) is the covariate vector and *h*_0_(*t*) is the baseline hazard function. CoxPH is a semi-parametric model and the baseline hazard function is canceled out when taking the ratio of two hazard functions.

The penalised Cox model is another extension of the CoxPH model that helps to prevent overfitting. The L1 regularized CoxPH model adds a scaled sum of absolute values of the magnitude of model coefficients, that is 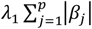 as the regularization term to the partial log-likelihood. Other regularizers can be used such as L2 regularization, that is 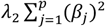, or other scaled sums of non-negative penalties of the *β*′*s*, such as in the following general penalised partial log-likelihood:

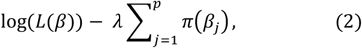

where *L*(*β*) is the partial likelihood as for example given in (Tibshirani, 1997; Equation 2) [8] and then optimization takes place[9] [10] [11]. Using the L1 penalty in Equation (2) gives the Lasso Cox estimation and using the L2 penalty gives the Ridge Cox solution, respectively. If instead of a single regularization term we consider a weighted average of the L1 and L2 penalty, we obtain the ElasticNet Cox model. One remarkable characteristic of the Lasso Cox model and the Elastic Net Cox model is that they can simultaneously perform feature selection and prediction, because some of the beta parameters can be penalised all the way to 0 when maximizing (2).

### 2.2 Modern machine learning models

Recent years have seen a strong surge in the use of modern ML methods in health as a result of their exceptional performance in many other areas, such as in finance [12], environment [13] and internet of things [14]. Notable examples in health include the application of Random Survival Forest (RSF) on complex metabolomics data [15] and SurvivalSVM to the survival of prostate cancer patients [16]. Both approaches are survival analysis extensions to two widely used ML algorithms for binary classification, namely Random Forest and SVM.

SurvivalSVM was developed by Van Belle et al [17] for time-to-event data. It is a variant of the regularized partial log-likelihood function (2) above but has a different penalty term. In contrast to using 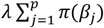, SurvivalSVM uses penalised splines and then applies both, ranking constraints and regression constraints to the corresponding partial log-likelihood function. SVM with those constraints enables models for high-dimensional omics data to have more flexible structure, e.g. additive (non)-linear models. One distinct feature of SurvivalSVM is that it treats the prognostic problem as a ranking problem and therefore, the estimation of the hazards is not directly incorporated in the model.

RSF was first proposed by [18] as an extension of Random Forest to model censored survival data. Random Forest [19] is a non-parametric bagging-based ensemble learning method that adds variation in the training datasets by bootstrapping the data. Multiple models are generated based on many resamples. The ensemble prediction result is then an average of these multiple models or the result of a majority vote. The key components in our application of RSF are that we use Harrell’s C-index to evaluate the survival tree instead of the mean square error for regression problems or confusion matrix for classification problems, and that we use the log-rank score in each node as the stopping rule.

Another ensemble-based approach is the boosting method, which contains multiple learners and sequentially gives more weight to weak learners to enhance predictability. For example, the Cox boosting model [20] [21] [22] [23] is developed based on Cox models with boosting being applied to the estimation of the regression parameter vector *β* in Equation (1). There are two popular approaches to update *β*: the first is the model-based approach that leads to the mboost method, the second is the likelihood-based approach that leads to the CoxBoost method (benchmarked in this study).

These models so far only focus on optimising a single objective. Because survival data is time dependent, it is natural to have multiple tasks related to one or more time points of interest. This naturally leads to multi-task learning, a method that deals with the need to predict for more than a single response variable, based on joint optimization of multiple likelihood functions corresponding to each task. The multi-task logistic regression model (MTLR) by Yu et al. [24] is a survival model for multiple time points, where for each, the task is to predict survival using a logistic regression model and the parameters from each model are estimated simultaneously in the maximization of the joint likelihood function.

More recently, the ML and artificial intelligence communities refer to the methods described above as classical ML methods due to the emergence of deep learning (DL), a conceptual advancement based on neural networks (NN). In survival analysis, a number of DL survival models were developed such as Cox-nnet [25], DeepSurv [26] and DeepHit [27]. The key concept here is having different loss functions that particularly target either the hazard or the survival probability for those neurons in hidden layers when building the DL architecture. High dimensional complex biological information can be better represented with the application of those hidden layers [28] and through relaxing the proportional hazard assumption.

### 2.3 Feature selection methods applied to survival models

The input features are fundamental to every statistical or machine learning model, and the survival model is no exception [29]. Wrapper and filter [30] are two feature selection methods that are widely used for not only regression and classification models, but also survival models.

The wrapper approach is a model-dependent method in which the performance of the model determines the selection of subsets of features. Stepwise feature selection approaches fall into this category since one feature is deemed to be included or deleted as the model’s performance improves. Other more computational approach such as the genetic algorithm (GA) [31], which was originally developed to solve an optimization problem, has been extended to use as a feature selection approach [32]. The main idea is to start with an initial set of features to then replace it with one that includes features from other parts of the data to optimize the classification accuracy based on a linear discrimination analysis model.

The filtering approach, on the other hand, is a model-independent feature selection method that produces a subset of features without involving the models. This step often occurs outside and before building prediction models. Many of these strategies select features using hypothesis testing statistics from a univariate study. With the advent of omics data in the 1990s, the statistics community embraced the development of differential expression (DE) analysis, which is a filter type feature selection method for identifying promising genes/features using “parallel univariate strategies” based on linear modelling [33].

### 2.4 Classical performance evaluation metric for survival data

Classically, survival analysis is evaluated in three broad settings: the concordance index, the Brier score and the time-dependent AUC. Similar to evaluating classification and regression models, metrics for calibration and discrimination frameworks are developed with incorporating censoring by applying rank-based methods or error based methods together with a weighting scheme.

#### 2.4.1 C-index and its extension in survival analysis

C-indices in survival analysis are concordance-based methods, where ‘concordance’ measures how close a prediction is to the truth. The first C-index for survival analysis was introduced by Frank. E. Harrell [34], as a time-independent performance measure. C-indices range from 0 to 1, where 1 means perfect performance and 0 means worst possible performance. If a model would not take into account any information from the data, that is a random prediction is made, then the corresponding C-index would be around 0.5. For most clinical datasets, a C-index around or larger than 0.6 is considered an acceptable prediction. Harrell’s C-index [35] defines concordance by looking at ranks of pairs of subjects in the data (there are n choose 2 pairs for data with n subjects). Harrell’s C index further depends on the censoring distribution of the data, is motivated by Kendall’s tau statistic and is closely related to Somers’ D. When ranking the subjects, censored subjects are excluded; and pairs included in the formula are only those comparable, non-censored pairs. There are different versions of the C-index, where the differences come from the different ways that censored subjects are ranked.

We will use the following three concordance indices: Begg’s C-index, Uno’s C-index and GH C-index. First, Begg’s C-index [36] uses KM estimation to incorporate both censored and uncensored subjects by assigning different weight to them. Second, Uno et al [37] develop a new way to calculate the rank with the help of inverse probability of censoring weight (IPCW). Third, the GH C-index [38] changes the concordance function into a probability function based on the Cox model estimation and then approximates its distribution which is robust to censoring.

#### 2.4.2 Brier score

The Brier score [39] [40] uses IPCW to handle censored subjects when measuring discrepancy between the estimated values and the actual values. This score can be considered as a similar measure to the mean squared error (MSE) in regression models to some extent. Like the MSE, the Brier score takes a value greater than 0 that depends on the data and the smaller the Brier score the better. However, to have better interpretability, the integrated Brier score (IBS) is introduced which also takes values between 0 and 1 - it averages the loss over time in situations where there is no interest in a particular time point but performance is with regards to all time points as a whole.

#### 2.4.3 Time-dependent AUC

The time-dependent AUC is inspired from binary classification model evaluations. The receiver operating characteristic (ROC) curve is a classical model assessment plot that examine the relationship between the sensitivity and the false positive rate. The area under the ROC curve is termed AUC (area under the curve). In survival analysis, event statuses are changing over the time, which requires a dynamic measurement to discriminate the predicted versus the actual. Chambless and Diao [41] were the first to propose a time-dependent AUC for survival analysis. They define the AUC(t) as the probability that a person with disease onset by time t has a higher score than the person with no event by timet. Changes of model predictability for different time points can therefore, be visualized by time-dependent AUC curves, which allows people to compare long time versus short time predictability.

## 3. Material and Methods

### 3.1 Datasets: six clinical and ten omics data sets

Clinical datasets - Six clinical datasets with different sample sizes and disease types are selected (see references in Supp Table 1).

- Veteran data is a survival dataset from the randomised trial of two treatment regimens for lung cancer obtained from the R package “survival”. There are 6 measured features in this data.
- Pbc data (5 clinical feature, 312 patients) from the Mayo Clinic trial in primary biliary cirrhosis (Pbc) of the liver conducted between 1974 and 1984; obtained from the R “RandomForestSRC”.
- Lung data (7 features, 228 patients) contains patient survival information with advanced lung cancer from the North Central Cancer Treatment Group and is available from the R package “survival.
- ANZ data (ANZDATA): Australia & New Zealand Dialysis and Transplant Registry data containing graft survival information and electronic clinical records for kidney transplantation recipients in Australia and New Zealand from 30^th^ June 2006 to 13^th^ November 2017. This data contains records for both living and deceased donors and also multi-organs transplants. We processed the raw data, restricting the transplant date to be after 2008-09-18 and retained deceased donor kidney transplants only. Missing records are excluded, resulting in 3323 patients and 38 features containing patient, donor and donor-recipient human leukocyte antigen (HLA) compatibility.
- UNOS_Kidney data: Organ transplant data based on the Organ Procurement and Transplantation Network (OPTN)-United Network for Organ Sharing (UNOS) in the US (based on OPTN data as of March, 2020). We selected a random sample of 3000 records associated with deceased donor kidney transplantation only with 99 features containing recipients, donors and donor-recipient HLA compatibility. Missing values are imputed using the R package “MICE”.
- Melanoma_clinical data, extracted from melanoma data [42] [43]: A in-house dataset collected as part of multi-omics study, is the part that contains clinical information for patients. After deleting all missing values, we have 88 patients with stage three melanoma disease measured by 14 clinical features.

Omics datasets - We consider eight published data and two in-house melanoma cancer datasets. A summary of the size and censoring rate of all datasets can be found in Supp Table 1.

- Two ovarian cancer gene expression datasets, downloaded from the R package “curatedOvarianData”. Curation and analysis pipeline of this data follow [44]. Ovarian1 is the “GSE49997_eset” data (194 patients/16047) genes, Ovarian2 is the “GSE30161_eset” data (58/19816).
- Another six gene expression datasets are available online from work by Yang and colleagues [45], namely GE_1, GE_2, …, GE_6. For GE_3, log2 transformation is applied, followed by a KNN imputation with 10 nearest points. For GE_6, median normalisation is applied. For others, no further pre-processing was performed.
- Melamona_itraq and Melanoma_nano are two in-house melanoma omics datasets, the first is a protein expression dataset from the iTRAQplatform and the second is a Nanostring dataset from the above melanoma study and pre-processing steps are described in the respective papers. The itraq protein expression data has 41 patients with 640 proteins. The nanostring data has 45 patients with 204 genes [46], and the GEO ID is “GSE156030”.

### 3.2 Benchmarking framework/procedure

#### Evaluation metrics

We examine model performance metrics that can be broadly grouped into four categories and assess performance in terms of each methods’ flexibility, predictability, stability and computational efficiency detailed in Supp Table 2 and briefly summarized as follows:

i. We measure *model flexibility* by looking at whether a given method can handle different data modality, different level of sparsity, represents multiple ways including the type of data required (clinical, omics), type of input required (categorical, numerical), sparsity of the data allowed (yes, no) and prediction ability evaluation metrics allowed.
ii. We measure *model predictability* using three different metrics: C-index, time-dependent AUC and Brier score. We apply four different modified versions of C-index: Harrell’s C-index, Begg’s C-index, Uno’s C-index and GH C-index. For identification of different time points, we equally divided the survival time ranging from the 1st quartile to the 3rd quartile into fifteen time points for each dataset, and therefore, we obtained fifteen AUC values corresponding to each time point. As for the Brier score, we calculated the raw Brier score and the IBS.
iii. We measure *model computational efficiency* using both computational time and memory. Computational time is calculated using the “Sys.time” function in R. Memory is calculated using the “Rprof” function in R and the total memory used is summarised for each experiment.
iv. We measure *model stability* using model reproducibility and the standard deviation (SD) of model predictability metrics. Model reproducibility is defined as the proportion of successful runs among all the runs attempted. For each model predictability metric, we calculated its SD. We then ranked the values for all the methods for each dataset from the most stable (smallest SD) to the least stable (largest SD).

#### Benchmarking methods

All methods evaluated are described in detail in the Supp Table 3. All compared methods (Supp Table 3) and evaluation metrics (Supp Table 2) are applied and evaluated on real-world datasets listed in Section 3.1. We apply 20 times (runs) repeated 5-fold cross validation using RStudio server with 15 cores in parallel. For each run, the whole data is split into a training dataset (80%) and a testing dataset (20%) with each method trained using the training dataset and values of evaluation metrics calculated using the testing dataset. Detail about the packages and parameters can be found in (Supp Table 3) and functions used to evaluate the methods are shown in (Supp Table 2).

## 4. Results

### 4.1 Comprehensive benchmarking framework

To comprehensively evaluate strength and weakness of the survival analysis approaches, we select 20 representative methods from our extensive literature review and study their performance when applied to 16 diverse datasets. The performance of each method is measured against 11 metrics representing multiple aspects, including feasibility, predictability, stability and computational efficiency. There are three key aspects of our comparison framework SurvBenchmark as depicted in Figure 1: (i) Practical focus through applying the framework to a broad range of datasets and by including a taxonomic methods system that evaluates multiple aspects; (ii) Extensive comparison of methods from classical to state of the art ML approaches; (iii) Comprehensive evaluation of the model performance with the utilisation of a customizable weighting framework.

**Figure 1:**
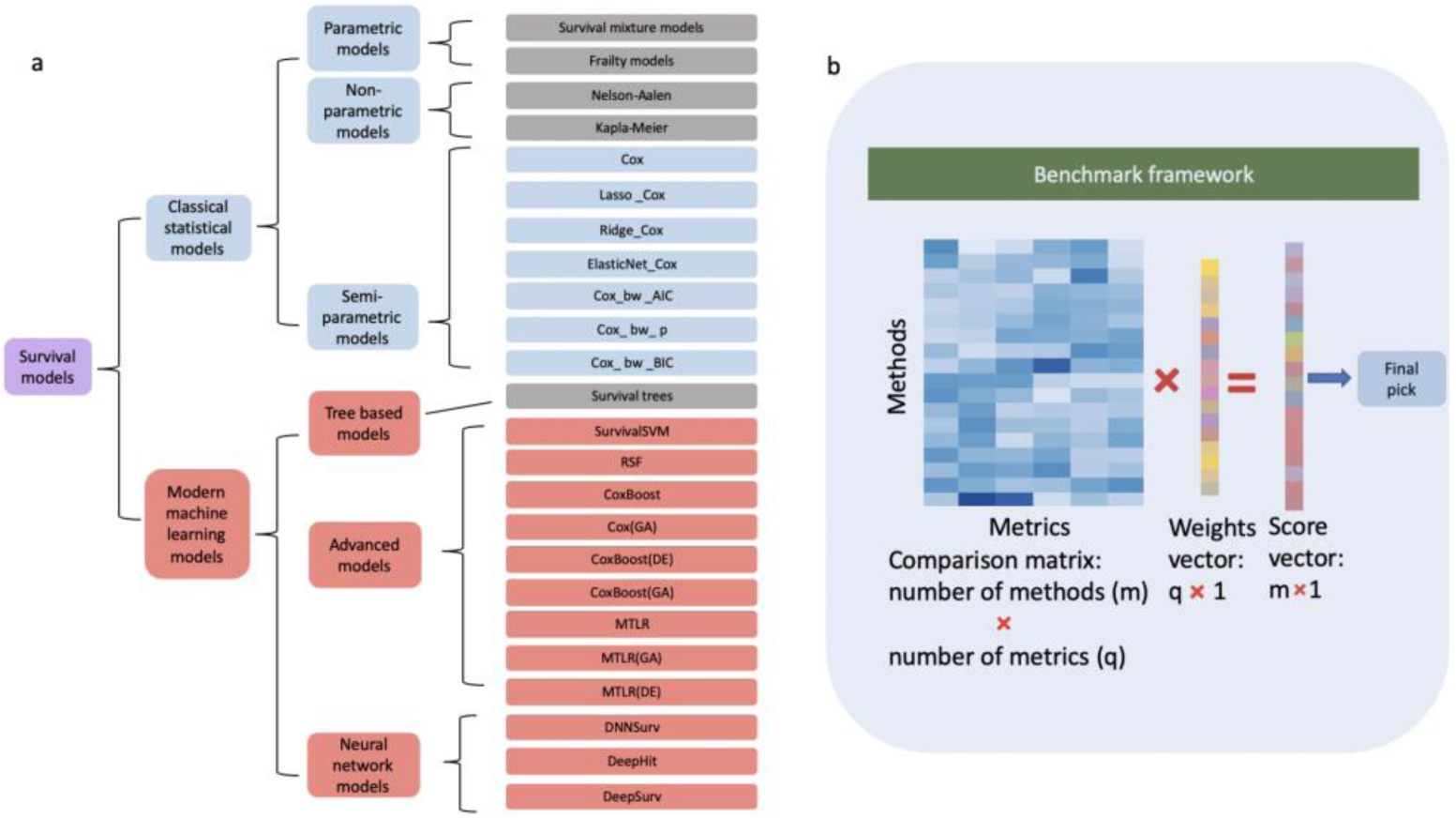
SurvBenchmark--Schematic view of our benchmark framework. (a) An overview of survival models. We broadly classify current models into two categories; classical statistical models (top group) and modern machine learning models (bottom group). Each of these categories can be further subdivided as presented in the hierarchical chart. All models in blue and red colored boxes are implemented in this current benchmark study. (b) A graphical representation of the SurvBenchmark framework. The methods and evaluation metrics are summarised in a matrix with a flexible user defined weights vector.

### 4.2 Practical consideration in assessing model performance

Many comparison studies define method performance solely in terms of method predictability, with only a few studies taking into account computational time. Often the feasibility of the method is not properly considered or discussed. Practically, it is paramount that a method can be applied to the data at hand, based on both the flexibility (data modality, sparsity) and computational requirement.

Given the diverse collection of data characteristics that is now available in the biomedical field, not all survival approaches are feasible to be applied to all data types. For example, some classical Cox models (Figure 2a, top left from column 1 to 10, row 1 to 4; a blue box indicates ‘method not feasible’), e.g. because it cannot handle large p (features) small n (samples) datasets (such as GE-1) which is a distinct feature of any molecular (omics) study. Advanced feature selection methods together with ML survival models such as CoxBoost(DE) can only take numerical data as the input (purple box for input type, where model characteristics are coded using 0, 1 and 2 with questions defined as below. Is input type numeric only? Yes: numerical only. No: both numerical and categorical are ok. Is output type survival risk? Yes: survival risk. No: survival probability. Can the model handle n<p situation? Yes: it can. No: it cannot. The other case: output is the rank of survival risk.).

**Figure 2:**
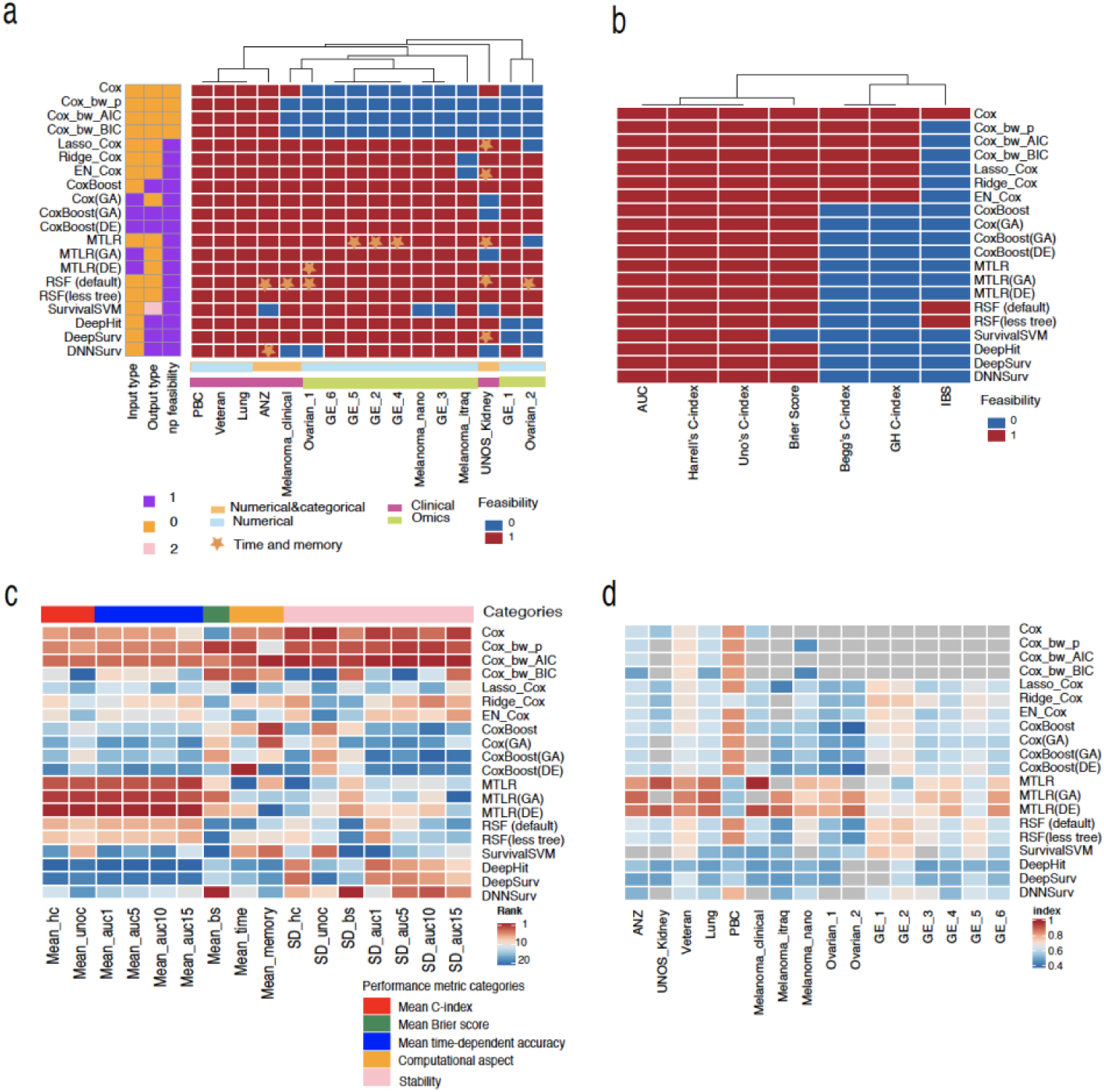
Summary heatmaps. (a) Summary for method flexibility and computational efficiency. (row: methods, column: datasets, feasible: red, not feasible: blue, feasible, star: memory and time consuming, model characteristics: yes, 1, no, 0, the other case, 2) (b) Prediction ability evaluation metric flexibility. (row: methods, column: prediction ability evaluation metrics, feasible: red, not feasible: blue) (c) Rank heatmap for method overall performance (row: methods, column: performance metrics, red to blue: top to bottom rank) (d) Harrell’s C-index heatmap (row: datasets, column: methods)

Next, we look at the computational aspect, and we notice that DL based methods are computationally inefficient as highlighted by the star icon (Figure 2a). From the many rowwise stars, we observe that RSF (5 stars) and MTLR (4 stars) are not as computationally efficient as Cox-based approaches such as Lasso_Cox (1 star) and CoxBoost (0 stars).

Lastly, a summary tabulating the feasibility associated with each of the evaluation metrics for prediction is provided in Figure 2b. The results highlight that Begg’s C-index and GH C-index are applicable only for Cox methods (red indicates feasibility), that the integrated Brier score can be calculated for Cox model and RSF (red), and that the Brier score cannot be calculated for SurvivalSVM (blue).

### 4.3 Performance evaluation from multiple perspectives: no ‘one size fits all’

To achieve a comprehensive overview of different survival approaches, we assess method performance from multiple perspectives across a large collection of datasets. Here, we color the methods according to their performances for all three broad categories: model predictability, model stability and computational efficiency (Figure 2c shows ranks of those methods where red means the best and blue the worst; similarly, Figure 2d shows Harrell’s C-index values with red referring to high values and blue to small values). We find that no method performs optimally across all three categories and there are various trade-offs among the categories.

For model predictability, we use seven different measures based on C-index, Brier score and time-dependent AUC. Here, MTLR-based approaches perform significantly better than others, which is most apparent by looking at the performance results using C-index and time-dependent AUC. In order to further examine whether MTLR-based approaches have similar performance across all datasets, we show our examination on one specific criteria (the most popular Harrell’s C-index; Mean_hc). In Figure 2d we demonstrate that MTLR has optimal performance for all but one of the six clinical datasets with PBC having optimal performance for one of the clinical datasets. Variants of MTLR (MTLR(GA) and MTLR(DE)) outperformed MTLR when applied to any of the ten omics datasets suggesting the performance of the approaches depend on the type of dataset.

For computational efficiency as measured by computational time and memory usage, the best performing methods are classical Cox-based models and CoxBoost. In particular, Cox, Cox_bw_AIC and Cox_bw_BIC are the top three performing methods for computational time (Figure 2c). For model stability, we have seven criteria and they are based on calculating the standard deviation (SD) of predictability metrics described above. Similar to the the computational efficiency performance, when using SD-criteria, Cox, Cox_bw_AIC and Cox_bw_BIC are also the top 3 performing methods in all but one criteria, the exception is the standard deviation of Brier score (SD_bs), where DNNSurv ranks first suggesting its ability to discriminate survival probabilities for different observations.

Despite having the best performance in both computational efficiency and stability for Cox-based methods, their predictability falls behind MTLR-based methods. On the other hand, MTLR-based methods clearly have the best predictability but they are not efficient and stable enough [47]. In conclusion, these observations demonstrate that no method performs optimally for all those categories.

### 4.4 Cox-based modern ML methods have similar prediction performance compared to classical Cox-based methods

To understand the gain in model predictability from Cox-based modern ML methods (CoxBoost, Coxboost (GA)), we compare these models with classical Cox-based methods (Lasso_Cox, EN_Cox) which are used as a gold standard method in many studies. Our results indicate that they have similar performance (Figure 3) across a large collection of datasets. For example, in the ANZ data, which is a representative clinical dataset, we observe similar model predictability measured by both Harrell’s C-index and Brier score. For GE_5, a representative dataset of omics with large p small n data characteristics, the same conclusion is drawn. This suggests the performance of modern ML methods in complex health and clinical data is not as clear cut as in some other domains.

**Figure 3:**
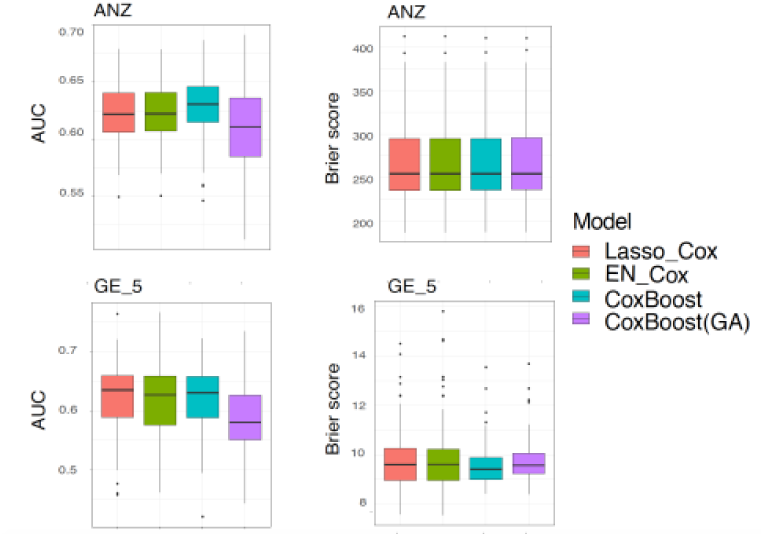
Prediction ability for cox-based methods. Top left: Harrell’s C-index on ANZ data. Top right: Brier score on ANZ data. Bottom left: Harrell’s C-index on GE_5. Bottom right: Brier score on GE_5.

### 4.5 Data dependent model performance for different time

To study the model performance over time, we visualize this using the time-dependent AUC curves for all methods. Here we observe among two representative clinical datasets (PBC, UNOS_Kidney) and two omics datasets (GE_2, GE_4) in Figure 4, not all curves are parallel to each other, indicating that the behaviour of model predictability for different time points is data dependent (see Supplementary Figure 1 for further results).

**Figure 4:**
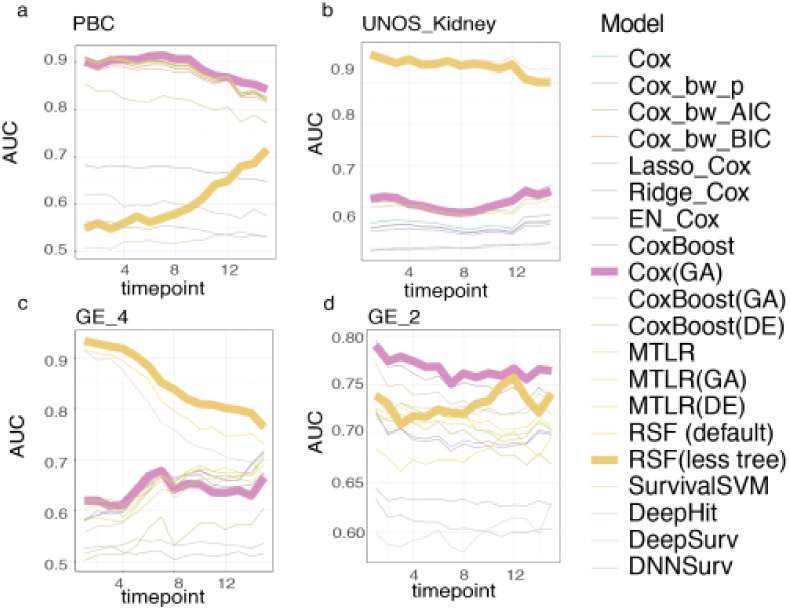
Time-dependent AUC curves for (a) PBC data; (b) UNOS_US data; (c) GE_4 data and (d): GE_2 data. Two selected models: Cox(GA), RSF (less tree)]

We pick two representative models (Cox(GA) & RSF(less tree)) to demonstrate this data dependent model behaviour. For UNOS_Kidney data and GE_2 data, the curves are approximately horizontal, which indicates the consistency of short-time, medium time and long-time model predictability. In contrast, for PBC data and GE_4 data, model predictability changes along those time points.

## 5. Discussion

This benchmark study comprehensively evaluated the relevance and usefulness of survival models in practice, where emphasis is on performance over diverse datasets. In our review we assessed a broad variety of survival methods from classical Cox-PH models to modern ML models. The findings of our systematic assessment will provide specific guidance for translational scientists and clinicians, as well as define areas of potential study in both survival methodology and benchmarking strategies.

In recent years, there is a clear shift in how survival data is analysed, from modelling directly the hazard function to building models directly on survival functions. Conceptually, modelling hazard functions is a good way to identify key risk factors related to various patients’ risk levels. On the other hand, if our key criterion is to predict accurately survival, modelling survival probability directly improves predictability. Methods including MTLR, DNNSurv and SurvivalSVM which directly model the survival function showed better performance in terms of model predictability, this is consistent with what [24] have commented on when discussing the performance of their proposed MTLR method.

It is striking that MTLR shows remarkable model predictability in our benchmark study. We now highlight technical advantages, disadvantages as well as its applications. Numerous reasons could contribute to the better model prediction performance of the MTLR-based approaches. These include the three main reasons as discussed by [24]: direct modeling of the survival function, simultaneous building of multiple logistic regression models, and dynamic modeling. Interestingly, the majority of extended MTLR models since 2011 are based on neural networks as researchers extend the concept to account for nonlinearity in datasets [48]. To date, only a limited number of studies have applied MTLR in health using clinical data in HIV patients [49] or on large omics datasets to predict patient survival in breast and kidney cancers [50]. Given its outstanding model predictability observed for most of the datasets in our study, we believe there is opportunity to use MTLR more widely for survival risk modelling in Health contexts.

Model predictability is one of the key metrics to assess survival studies with Harrell’s C-index being currently the most popular. As this kind of ranking based concordance measurement is suitable to evaluate predicted outcomes with censored data, various concordance indices are developed using different methods to handle censoring such as Uno’s C-index using IPCW. Besides concordance indices, other predictability metrics such as the time-dependent AUC, which applies a similar idea as the AUC in binary classification but divides the whole time interval into multiple time points, are also adopted in some survival studies [51]. Given that model predictability could be measured by multiple types of indices, we suggest that hybrid evaluation metrics should be applied in practice to provide relatively comprehensive assessments for the fitted model.

Though many survival approaches are applicable to both clinical and omics data, there are a number of recently developed approaches that are specifically tailored for high dimensional omics data, such as CoxBoost. The rationale behind developing data-specific methods is to better capture the distinct data characteristics in either the clinical or omics studies. Clinical data usually include mixed modality variables, large sample sizes but have large n (observations) and small p (features). In contrast, omics data naturally comes a large collection of molecular features and with small n but their data type is homogenous. When it comes to various real-world datasets, performances are also affected by many other aspects besides data type (clinical, omics) such as data modality and therefore, it is challenging to directly examine whether those tailored methods indeed improve the performance.

Unlike in other approaches, there are some reproducibility concerns for the method DNNSurv in our empirical work: It failed to run for some of the cross-validation runs. Specifically, among all 100 runs, DNNSurv had a 100% completion rate for 5 out of the 12 applicable datasets (Supplementary Figure 2) only. For the remaining 7 datasets, completion rate was around 80% and as low as 63% for the Melanoma_itraq data. This instability is likely due tuning parameter sensitivity when sample size is small [52].

## Acknowledgements

The authors thank all their colleagues, particularly at The University of Sydney, Sydney Precision Bioinformatics Alliance and Charles Perkins Centre for their support and intellectual engagement.

## Data and code availability

For the ANZDATA, data request can be made through the ANDATA registry, and access to the data source will require HREC approvals. For the UNOS_kidney data, it can be requested from https://optn.transplant.hrsa.gov/data/. Codes for running those methods and evaluation measurements for an example dataset is available at https://github.com/SydneyBioX/SurvBenchmark.

## Funding

The following sources of funding for each author, and for the manuscript preparation, are gratefully acknowledged: Australian Research Council Discovery Project grant (DP170100654) to JYHY and SM, Australian Research Council Discovery Project grant (DP210100521) to SM, AIR@innoHK programme of the Innovation and Technology Commission of Hong Kong to JYHY. Research Training Program Tuition Fee Offset and Stipend Scholarship and the Dean’s International Postgraduate Research Scholarship (DIPRS) to YZ. The funding source had no role in the study design; in the collection, analysis, and interpretation of data, in the writing of the manuscript, and in the decision to submit the manuscript for publication.

## Conflict of Interest

none declared.

## Notes

https://github.com/SydneyBioX/SurvBenchmark

### Competing Interest Statement

The authors have declared no competing interest.

### Summary of Updates

Figure 1 and Figure 2 update to a clearer version; Supplemental files update

