## Supplementary Material for "SurvBenchmark: comprehensive benchmarking study of survival analysis methods using both omics data and clinical data"

**Supplementary Figures**

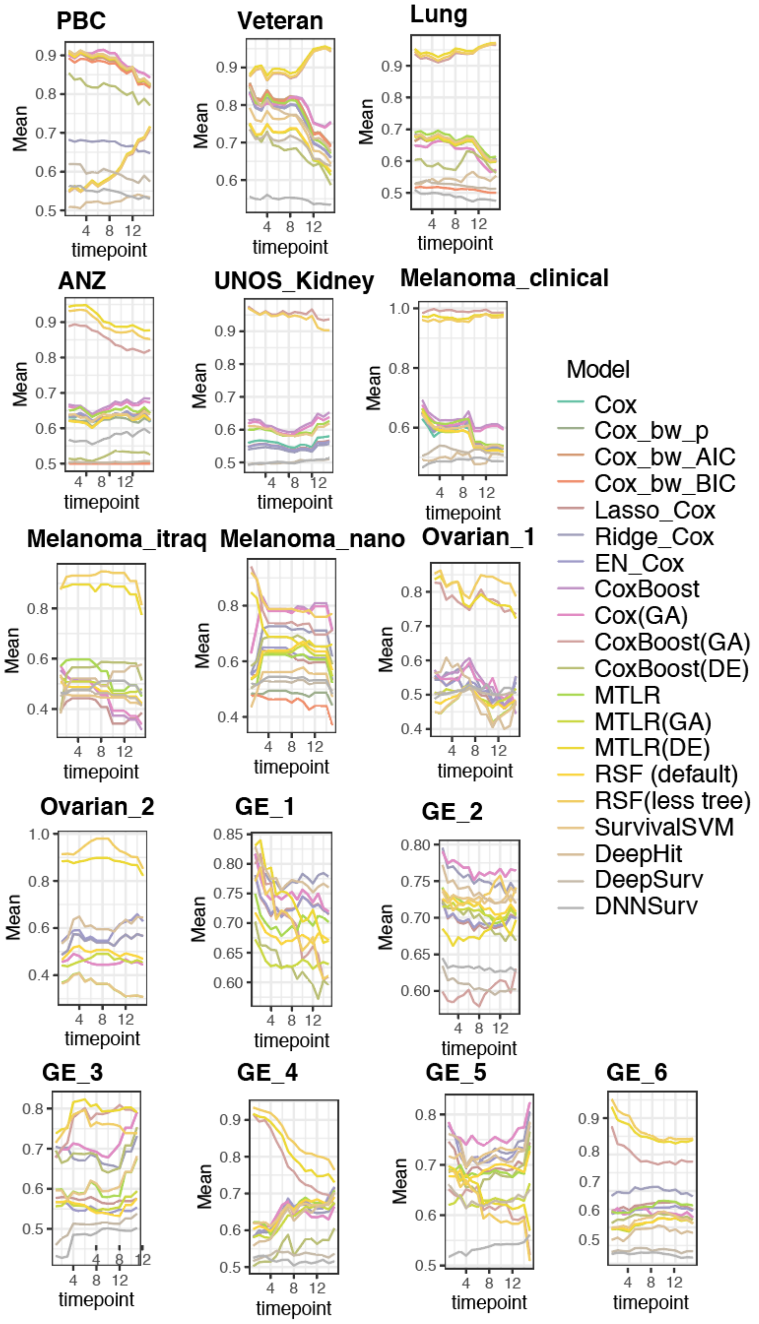


**Supplementary Figure 1: Data dependent performance for short and long time prediction.** We show the time-dependent model predictability (time-dependent AUC) for all models for each dataset across 15 time points. Different trends for those lines for different datasets indicate that model predictability for short-time versus long-time depends on the dataset.

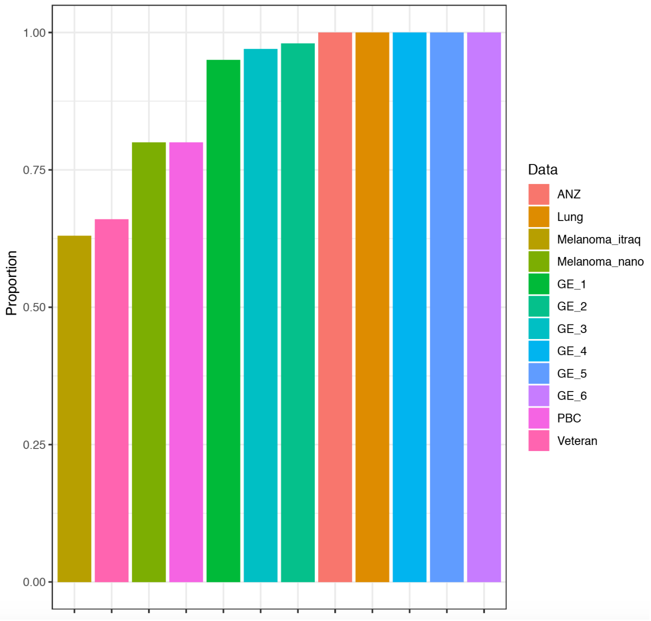


**Supplementary Figure 2: DNNSurv reproducibility.**
We show that DNNSurv has a 100% completion rate for 5 (ANZ, Lung, GE_4, GE_5, GE_6) out of the 12 applicable datasets. For the remaining 7 datasets, the completion rate is around 80% and completion rate was as low as 63% for the Melanoma_itraq data.

**Supp Table1.** Data table showing the names of datasets used in this paper in the first column. Datasets are ordered by the number of observations (second column, from smallest to largest). Censoring rate is rounded to 4 decimal places.

| Datasets summary | | | | | |
|---|---|---|---|---|---|
| **Dataset (name used in this paper)** | **Number of observations** | **No. of variables** | **Type of data** | **Censoring rate (rounded to 4 decimal places)** | **Reference** |
| Melanoma_itraq | 41 | 642 | Omics | 0.4146 | Wang,K.Y.X. et al. Cross-Platform Omics Prediction procedure: a game changer for implementing precision medicine in patients with stage-III melanoma.bioRxiv 2020.12.09.415927; doi: https://doi.org/10.1101/2020.12.09.415927 |
| Melanoma_nano | 45 | 206 | Omics | 0.4222 | Wang,K.Y.X. et al. Cross-Platform Omics Prediction procedure: a game changer for implementing precision medicine in patients with stage-III melanoma.bioRxiv 2020.12.09.415927; doi: https://doi.org/10.1101/2020.12.09.415927 |
| Ovarian_2 | 58 | 19818 | Omics | 0.3793 | Ganzfried,B.F. et al. (2013) curatedOvarianData: clinically annotated data for the ovarian cancer transcriptome. Database, 2013. |
| GE_5 | 78 | 4753 | Omics | 0.5641 | van 't Veer,L.J. et al. (2002) Gene expression profiling predicts clinical outcome of breast cancer. Nature, 415, 530–536. |
| GE_3 | 86 | 6288 | Omics | 0.7209 | Bullinger,L. et al. (2004) Use of Gene-Expression Profiling to Identify Prognostic Subclasses in Adult Acute Myeloid Leukemia. New England Journal of Medicine, 350, 1605–1616. |
| Melanoma_clinical | 88 | 16 | Clinical | 0.3939 | Wang,K.Y.X. et al. Cross-Platform Omics Prediction procedure: a game changer for implementing precision medicine in patients with stage-III melanoma.bioRxiv 2020.12.09.415927; doi: https://doi.org/10.1101/2020.12.09.415927. |
| GE_1 | 115 | 551 | Omics | 0.6670 | Sorlie,T. et al. (2003) Repeated observation of breast tumor subtypes in independent gene expression data sets. Proc. Natl. Acad. Sci. U. S. A., 100, 8418–8423. |
| GE-_4 | 116 | 4753 | Omics | 0.5641 | van de Vijver,M.J. et al. (2002) A gene-expression signature as a predictor of survival in breast cancer. N. Engl. J. Med., 347, 1999–2009. |
| Veteran | 137 | 8 | Clinical | 0.0657 | Kalbfleisch,J.D. and Prentice,R.L. (2002) The Statistical Analysis of Failure Time Data. Wiley Series in Probability and Statistics. |
| Ovarian_1 | 194 | 16050 | Omics | 0.7062 | Ganzfried,B.F. et al. (2013) curatedOvarianData: clinically annotated data for the ovarian cancer transcriptome. Database, 2013. |
| Lung | 228 | 9 | Clinical | 0.2763 | Loprinzi,C.L. et al. (1994) Prospective evaluation of prognostic variables from patient-completed questionnaires. North Central Cancer Treatment Group. J. Clin. Oncol., 12, 601–607. |
| GE_6 | 240 | 7401 | Omics | 0.4250 | Van Houwelingen,H.C. (2004) The Elements of Statistical Learning, Data Mining, Inference, and Prediction. Trevor Hastie, Robert Tibshirani and Jerome Friedman, Springer, New York, 2001. No. of pages: xvi 533. ISBN 0-387-95284-5. Statistics in Medicine, 23, 528–529. |
| GE_2 | 295 | 4921 | Omics | 0.7322 | Beer,D.G. et al. (2002) Gene-expression profiles predict survival of patients with lung adenocarcinoma. Nat. Med., 8, 816–824. |
| PBC | 312 | 7 | Clinical | 0.5994 | Fleming,T.R. and Harrington,D.P. (2005) Counting Processes and Survival Analysis. Wiley Series in Probability and Statistics. |
| UNOS_Kidney | 3000 | 101 | Clinical | 0.7350 | OPTN data |
| ANZ | 3323 | 40 | Clinical | 0.8739 | ANZDATA |

**Supp Table 2.** Different evaluation criteria for assessing the performance models

| **Evaluation criteria** | | |
|---|---|---|
| **Class** | **Name** | **Description** |
| **Model flexibility** | Type of data required | Describes the source of the data obtained, either clinical data or omics data. |
| | Data input | Describes the different types of data modality such as categorical, numerical or mixture. |
| | Data sparsity | Binary yes or no measure whether the model can handle a high level of sparsity in the data. |
| | Prediction ability evaluation metrics allowed | A certain evaluation metric can only be applied to a specific type of survival model. This is a 2-dim model cross predictability evaluation metric summary and whether a model can be assessed by a metric is recorded with yes or no values. |
| **Model predictability** | Harrell's C-index | Harrell's method to calculate the C-index. This is applied to all methods using the R function "rcorr.cens" in package "Hmisc". This value ranges from 0 to 1 with 0.5 representing random guesses and the higher the value the better the concordance, i.e. model predictability. |
| | Begg's C-index | Begg's method to calculate the C-index. This is applied to applicable methods using the R function "BeggC" in package "survAUC". |
| | Uno's C-index | Uno's method to calculate the C-index. This is applied to all methods using the R function "UnoC" in package "survAUC". |
| | GH C-index | Gonen and Heller's method to calculate the C-index. This is applied to applicable methods using the R function "GHCI" in package "survAUC". |
| | Time-dependent AUC for time t | This is the Chambless and Diao estimator of cumulative/dynamic AUC for right-censored time-to-event data for time t. This is applied to all methods using the R function "AUC.cd" in package "survAUC". The interpretation of this AUC value for a specific time t is the same as AUC in classification models. |
| | Brier score | This calculates the Brier score for all methods using the R function "pec" in package "pec". The smaller the value, the better the prediction. |
| | Integrated Brier score | This gives the scaled version of the Brier score. This value is between 0 and 1 (the smaller the value, the better the prediction) and for the constant prediction probability 0.5, the value is 0.25. This is applied to feasible methods using the R function"crps" in package "pec". |
| **Computational efficiency** | Computational time | Represents how long it takes the method to run for each dataset. This is calculated using the "Sys.time" function in R. |
| | Total Memory | Represents how much memory is required to run the method for each dataset. This is calculated using the "Rprof" function in R which returns the memory consumed for each sub function and then we calculate the total memory which is the sum of all of those. |
| **Model stability** | Reproducibility | Measures the proportions of successful runs among all those 100 runs attempted. Some methods are not fully successful for all datasets, such as DNNSurv and the corresponding successful proportions are plotted. (Supplementary Figure 2) |
| | SD of model predictability metrics | Measures the standard deviation (SD) of each model predictability metric. Values are ranked from 1 (smallest SD) to 20 (largest SD) for all those 20 methods within each dataset. |

**Supp Table 3. Details of the methods used in this study**

| Method name | Method name in this paper | R function name | R package name | Data type | Parameters | Input type | Output type | N<p feasibility |
|---|---|---|---|---|---|---|---|---|
| **Cox** | Cox | coxph | survival | Clinical | NA | Categorical/numerical | risk | no |
| **Cox with baclward elimination using AIC** | Cox_bw_AIC | cph, fastbw | rms | Clinical | rule="aic", sls=.05,k.aic=2 | Categorical/numerical | risk | no |
| **Cox with baclward elimination using p value** | Cox_bw_p | cph, fastbw | rms | Clinical | rule="p", sls=.05 | Categorical/numerical | risk | no |
| **Cox with baclward elimination using BIC** | Cox_bw_BIC | cph, fastbw | rms | Clinical | rule="aic",sls=.05,k.aic = log(as.numeric(table(train$status)[2])) | Categorical/numerical | risk | no |
| **Lasso cox** | Lasso_Cox | penalized | penalized | Clinical | Lambda1=1, lambda2=0 | Categorical/numerical | Survival probability | no |
| **Ridge cox** | Ridge_Cox | penalized | penalized | Clinical | Lambda1=0, lambda2=1 | Categorical/numerical | Survival probability | no |
| **Elastic net cox** | EN_Cox | penalized | penalized | Clinical | Lambda1=1, lambda2=1 | Categorical/numerical | Survival probability | no |
| **Lasso cox** | Lasso_Cox | glmnet | glmnet | Omics | alpha=1, nfolds = 5,type.measure = "C" | Numerical | risk | yes |
| **Ridge cox** | Ridge_Cox | glmnet | glmnet | Omics | alpha=0, nfolds = 5,type.measure = "C" | Numerical | risk | yes |
| **Elastic net cox** | EN_Cox | glmnet | glmnet | Omics | alpha=0.5, nfolds = 5,type.measure = "C" | Numerical | risk | yes |
| **Random survival forest version1** | RSF(default) | rfsrc | RandomSurvivalForest | Clinical & omics | Default (ntree = 1000 , mtry = 10) | Categorical/numerical | risk | yes |
| **Random survival forest version2** | RSF(less tree) | rfsrc | RandomSurvivalForest | Clinical & omics | ntree = 100, mtry = 20 | Categorical/numerical | risk | yes |
| **Multi task logistic regression method** | MTLR | mtlr | MTLR | Clinical & omics | C1_vec = c(0.01,0.05,0.1,0.5,1,10) | Categorical/numerical | risk | yes |
| **DNNSurv (Deep learning survival model)** | DNNSurv | multiple functions as in Github codes | DNNSurv (https://github.com/lilizhaoUM/DNNSurv) | Clinical & omics | Default | Categorical/numerical | Survival probability | yes |
| **Boosting cox model** | CoxBoost | coxboost | CoxBoost | Clinical & omics | stepnumber=10,penaltynumber=100 | Categorical/numerical | Survival probability | yes |
| **Cox model with genetic algorithm as feature selection method** | Cox(GA) | GenAlg | GenAlgo | Omics | n.features=10 (for omics) , n.features=4 (for clinical) , generation_num=20 | Numerical | risk | yes |
| **Multi task logistic regression model with genetic algorithm as feature selection method** | MTLR(GA) | GenAlg | GenAlgo | Omics | n.features=10 (for omics) , n.features=4 (for clinical) ,generation_num=20 | Numerical | risk | yes |
| **Boosting cox model with genetic algorithm as feature selection method** | CoxBoost(GA) | GenAlg | GenAlgo | Omics | n.features=10 (for omics) , n.features=4 (for clinical) , generation_num=20 | Numerical | Survival probability | yes |
| **Multi task logistic regression model with ranking based method as feature selection meth** | MTLR(DE) | lmFit,eBayes | limma | Omics | n.features=10 (for omics) , n.features=4 (for clinical) | Numerical | risk | yes |
| **Boosting cox model with ranking based method as feature selection method** | CoxBoost(DE) | lmFit,eBayes | limma | Omics | n.features=10 (for omics) , n.features=4 (for clinical) | Numerical | Survival probability | yes |
| **Survival support vector machine** | SurvivalSVM | survivalsvm | survivalsvm | Clinical & omics | Default | Categorical/numerical | Risk rank | yes |
| **DeepSurv (Deep learning survival model)** | DeepSurv | deepsurv | survivalmodels | Clinical & omics | default: frac = 0.3, activation = "relu",<br>num_nodes = c(4L, 8L, 4L, 2L), dropout = 0.1, early_stopping = TRUE, epochs = 100L,<br>batch_size = 32L | Categorical/numerical | Survival probability | yes |
| **DeepHit (Deep learning survival model)** | DeepHit | deephit | survivalmodels | Clinical & omics | default: frac = 0.3, activation = "relu",<br>num_nodes = c(4L, 8L, 4L, 2L), dropout = 0.1, early_stopping = TRUE, epochs = 100L,<br>batch_size = 32L | Categorical/numerical | Survival probability | yes |
